# The multi-speed genome of *Fusarium oxysporum* reveals association of histone modifications with sequence divergence and footprints of past horizontal chromosome transfer events

**DOI:** 10.1101/465070

**Authors:** Like Fokkens, Shermineh Shahi, Lanelle R. Connolly, Remco Stam, Sarah M. Schmidt, Kristina M. Smith, Michael Freitag, Martijn Rep

**Affiliations:** Molecular Plant Pathology, University of Amsterdam, The Netherlands; Department of Biochemistry and Biophysics, Oregon State University, United States of America; Chair of Phytopathology, School of Life Sciences Weihenstephan, Technische Universität München, Germany; Institute for Molecular Physiology, Heinrich-Heine-Universität Düsseldorf, Germany; Departement of Biology, Oregon State University-Cascades, United States of America

## Abstract

*Fusarium oxysporum* is an economically important pathogen causing wilting or rotting disease symptoms in a large number of crops. It is proposed to have a structured, “two-speed” genome: i.e. regions containing genes involved in pathogenicity cluster with transposons on separate accessory chromosomes. This is hypothesized to enhance evolvability. Given the continuum of adaptation of all the genes encoded in a genome, however, one would expect a more complex genome structure. By comparing the genome of reference strain Fol4287 to those of 58 other *Fusarium oxysporum* strains, we found that some Fol4287 accessory chromosomes are lineage-specific, while others occur in multiple lineages with very high sequence similarity - but only in strains that infect the same host as Fol4287. This indicates that horizontal chromosome transfer has been instrumental in past host-switches. Unexpectedly, we found that the sequence of the three smallest core chromosomes (Chr. 11, 12 and 13) is more divergent than that of the other core chromosomes. Moreover, these chromosomes are enriched in genes involved in metabolism and transport and genes that are differentially regulated during infection. Interestingly, these chromosomes are –like the accessory chromosomes– marked by histone H3 lysine 27 trimethylation (H3K27me3) and depleted in histone H3 lysine 4 dimethylation (H3K4me2). Detailed genomic analyses revealed a complex, “multi-speed genome” structure in *Fusarium oxysporum*. We found a strong association of H3K27me3 with elevated levels of sequence divergence that is independent of the presence of repetitive elements. This provides new leads into how clustering of genes evolving at similar rates could increase evolvability.

**Author summary:** Fungi that cause disease on plants are an increasingly important threat to food security. New fungal diseases emerge regularly. The agricultural industry makes large investments to breed crops that are resistant to fungal infections, yet rapid adaptation enables fungal pathogens to overcome this resistance within a few years or decades. It has been proposed that genome ‘compartmentalization’ of plant pathogenic fungi, in which infection-related genes are clustered with transposable elements (or ‘jumping genes’) into separate, fast-evolving regions, enhances their adaptivity. Here, we aimed to shed light on the possible interplay between genome organization and adaptation. We measured differences in sequence divergence and dispensability between and within individual chromosomes of the important plant pathogen *Fusarium oxysporum*. Based on these differences we defined four distinct chromosomal categories. We then mapped histone modifications and gene expression levels under different conditions for these four categories. We found a ‘division of labor’ between chromosomes, where some are ‘pathogenicity chromosomes’ - specialized towards infection of a specific host, while others are enriched in genes involved in more generic infection-related processes. Moreover, we confirmed that horizontal transfer of pathogenicity chromosomes likely plays an important role in gain of pathogenicity. Finally, we found that a specific histone modification is associated with increased sequence divergence.

## Introduction

Genes encode products that are involved in a myriad of biological processes and are subjected to a wide range of selection pressures that may change over time. In pathogenic micro-organisms, fast-evolving genes are often clustered with transposable elements into separate regions, an observation that has led to the “two-speed genome” hypothesis concept [1-3]. A possible advantage for the separate genome regions with different evolutionary rates is that it might improve a genome’s ability to accommodate conflicting evolutionary demands, thus allowing an organism to adapt more quickly to new environments [1-4]. Pathogens secrete specialized proteins, called effectors, that facilitate colonization of a host. These effectors are, however, sometimes recognized by the host immune system and then trigger an immune response. Effectors that trigger an immune response are called avirulence factors. A pathogen can escape this recognition when the avirulence gene is mutated or no longer expressed during infection. There is accumulating evidence that this is more likely to occur when an avirulence gene is located near transposon sequences. The occurrence of multiple almost identical sequences in a genome increases the likelihood of homologous recombination that can result in genome rearrangements or deletions [5], which can contribute to pathogen adaptation [1-3,6-15]. In addition, transposon insertion into an avirulence gene can result in regain of virulence [16-19]. If the avirulence gene is indispensable for infection the only option to escape recognition is to alter the amino acid sequence of the effector [20-22]. Moreover, an effector needs to coevolve with its target in the host (see e.g. [23]). Effector genes and other genes that are located in ‘fast’ regions have been found to be under positive selection [24,25] and in general have more polymorphisms [26]. However, with the notable exception of repeat-induced point mutation (RIP) in *Leptosphaeria* species [27,28], no mechanism explaining the link between the proximity of transposons and elevated levels of sequence evolution has yet been firmly established.

Core and accessory chromosomes in fungi typically not only differ in terms of dispensability. They also show differences in gene- and repeat density and -in species that undergo a sexual cycle- in recombination frequencies [29,30] (see e.g. [13,31] for review). Recent studies on the chromatin of core and accessory regions showed that accessory regions are marked by histone H3 lysine 27 trimethylation (H3K27me3), sometimes called ‘facultative heterochromatin’ [31-34]. One hypothesis is that this allows for concerted regulation of genes involved in similar processes, such as infection [35].

As a basis for investigating the underlying mechanisms of genome compartmentalization and differences in evolutionary ‘speeds’, a detailed mapping of evolutionary ‘volatility’ across a genome is required and correlations with transposon proximity, histone modifications and gene expression must be established. In this study, we used 58 *Fusarium oxysporum* genome sequences for a detailed comparison of differences in dispensability and levels of sequence divergence between and within chromosomes. We combined this with functional annotation and gene expression data to study the extent to which evolutionary volatility is linked to involvement in infection. To study whether there are differences in chromatin-mediated regulation between and within chromosomes in Fol4287, and whether these differences correlate with differences in dispensability and sequence divergence and gene expression, we determined the distribution of histone marks associated with euchromatin (H3K4me2) and facultative heterochromatin (H3K27me3) in vitro and correlated these to gene annotation categories as well as levels of gene expression *in vitro* and *in planta*.

The *Fusarium oxysporum* species complex (FOSC) comprises strains that can infect over a 100 economically important crops, though most strains are not known to be pathogenic [36,37]. Individual pathogenic strains are mostly specialized towards a single host and are grouped according to host-preference into *formae speciales* (ff.spp.) [36,38-40]. Strains that belong to the same *forma specialis* (f.sp.) have similar effector repertoires [40]. In 2010, Ma et al. reported the genome sequence of the tomato-infecting strain *Fusarium oxysporum* f. sp. *lycopersici* 4287 (Fol4287). Out of 15 chromosomes in Fol4287, only 11 are largely syntenic with chromosomes of *Fusarium verticillioides*, a sister species that is estimated to have diverged about 11 Mya [37,41]. From this comparison, these 11 chromosomes constitute the core genome of Fol4287, whereas 4 chromosomes (3, 6, 14 and 15) and two large, presumable translocated, regions (chromosome 1b and 2b) are absent in *Fusarium verticillioides* and thus constitute the accessory genome of Fol4287. These chromosomes also have been reported to be largely absent in the strains *Fom*-5190a, *Foc*-38-1, *Fop*-37622, *Fo5176*, and *Fom001*, infecting *Medicago* spp., chickpea, pea, *Brassica* spp. and melon respectively [42], indicating that they are conditionally dispensable in the FOSC. Remarkably, strains that have lost ‘core’ chromosome 12 showed no reduction in virulence in experimental conditions and reduction of growth on only a few tested carbon sources. This indicates that chromosome 12 is also conditionally dispensable and thus not a core chromosome *sensu strictu* [43].

Interestingly, host-preference in FOSC is often polyphyletic [44-56]. Several studies revealed putative horizontal transfer events of effector genes between FOSC strains that infect the same host [40,57-59], indicating that horizontal transfer may play an important role in host switches [39,40,56,60,61]. Of the four accessory chromosomes in Fol4287, chromosome 14 is undoubtedly a pathogenicity chromosome: all fourteen SIX (Secreted In Xylem) genes – known effector genes in tomato-infecting *Fusarium oxysporum* – are located on chromosome 14 [62] (and our unpublished observations). Loss of chromosome 14 leads to loss of pathogenicity in Fol4287 [43]. Importantly, horizontal transfer experiments have shown that acquisition of chromosome 14 by the non-pathogenic strain Fo47 is sufficient to turn this strain pathogenic on tomato, albeit less virulent than the original donor [41,63]. The first horizontal transfer experiment reported for the FOSC used the tomato-infecting strain Fol007, which belongs to the same clonal lineage as Fol4287 yet has a different karyotype: Fol007 has a small chromosome that is not present in Fol4287. Interestingly, when this small chromosome co-transferred with chromosome 14, the recipient strains were much more virulent [41]. Recently, a similar experiment with the pathogenicity chromosome of Forc016, causing root-rot in cucurbits, shows that host-switches by horizontal chromosome transfer are not unique to tomato-infecting strains [64].

The polyphyletic origins of host-specificity, the fact that strains that infect the same host have similar effector repertoires, the frequent observations of horizontal transfer of effector genes, and the fact that horizontal transfer of a pathogenicity chromosome can render a non-pathogenic strain pathogenic all indicate that horizontal chromosome transfer causes host-switches in FOSC. Our genome dataset includes tomato-infecting strains that belong to different lineages, hence we also used it to detect putative recent horizontal chromosome transfer events.

## Results

### The accessory genome of Fol4287 consists of pathogenicity-related regions and lineage-specific regions

When compared to *F. verticillioides*, the accessory genome of Fol4287 consists of four chromosomes and two large segments on core chromosomes. We compared the genome of Fol4287 to that of a closely related strain that is pathogenic on melon, Fom001, and found that the core chromosomes are almost 100% identical between these strains, but that some of the accessory chromosomes are specific to Fol4287 (Fig S1). Chromosomal regions 1b and 2b and chromosome 15, however, align with the genome of the Fom001 with similar levels of sequence similarity as the core chromosomes (Fig S1). This suggests that host-preference is largely determined by genes that are located on three accessory chromosomes. We predict that genes on these chromosomes that are important for tomato infection are also present in other strains that are pathogenic on tomato.

To further assess which accessory regions correlate with phylogenetic clades (lineage-specific chromosomes), and which with *forma specialis* (pathogenicity chromosomes), we aligned 58 FOSC genomes to the genome of Fol4287 (Fig S2, Table S1). We found that the four accessory chromosomes are largely clade-specific. Alignments with genomes of strains outside the lineage of Fol4287 (including Fom001) mostly span less than 5 kilobases (kb) and have less than 92% sequence identity. Notable exceptions are chromosome 14 and a recently duplicated region on chromosome 3 and 6. These regions are present in more distantly related tomato-infecting strains with ∼100% sequence identity, compared to ∼98% identity between core chromosomes (Fig1A, Fig 2, Fig S2). For these regions synteny is also relatively conserved in tomato-infecting isolates, with aligned segments spanning up to 40 kb, whereas for the rest of the accessory genome this is between 5 and 10 kb (Fig S3).

**Fig 1.**
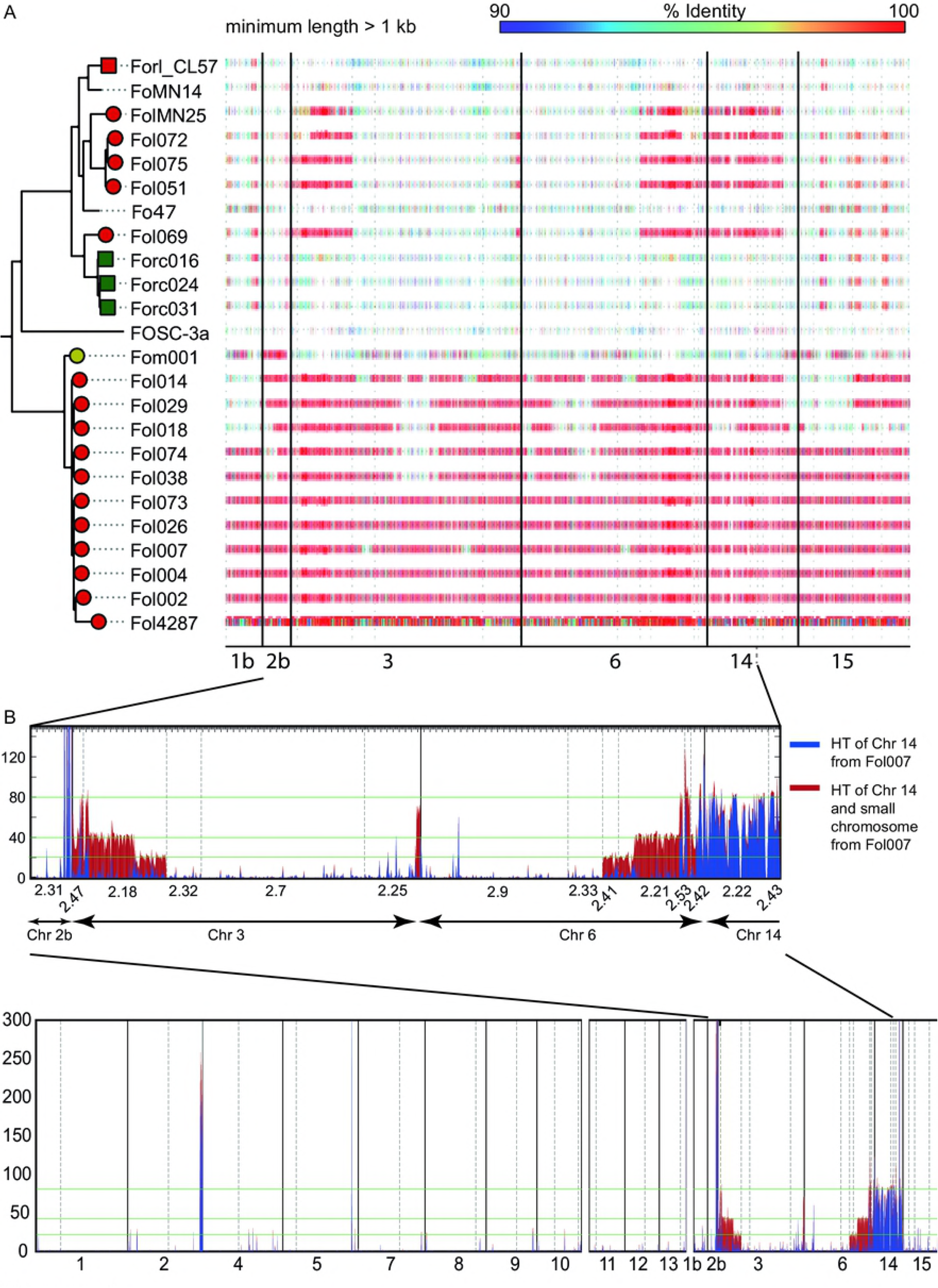
Lineage-specific and pathogenicity regions in the Fol4287 accessory genome. **A**. Horizontal bars -colored according to the percent identity in the alignment-indicate presence of Fol4287 accessory regions 1b and 2b and accessory chromosomes 3, 6, 14 and 15 in 23 other *Fusarium oxysporum* isolates. Leaf nodes in the phylogenetic tree are colored according to *forma specialis*, isolates causing wilting symptoms are represented with a circle, those that cause root rot are represented with squares and those that are non-pathogenic do not have a shape or color. Only alignments that span more than 1 kb and are more than 90% identical are included. Chromosomal regions 1b (Supercontig 2.27) and 2b (Supercontig 2.31), part of chromosome 3, part of chromosome 6 and chromosome 15 are mosty absent or present with relatively low sequence similarity outside the clonal line of Fol4287. In contrast, chromosome 14, part of chromosome 3 and part of chromosome 6 are present with ∼100% sequence identity in all tomato-infecting isolates that we queried. **B**. Two strains that respectively received one and two chromosomes in a horizontal transfer experiment involving Fol007 as chromosome donor and Fo47 as recipient, had been sequenced previously [62]. We mapped sequencing reads obtained from these strains on the genome of Fol4287. The density of mapped reads from the strain that received chromosome 14 is depicted in blue. The density of mapped reads from the strain that received a small chromosome in addition to chromosome 14 and is more virulent, is depicted in red. These read densities reveal that that the pathogenicity regions on chromosome 3 and 6 correspond to a small pathogenicity chromosome in Fol007. Read densities in regions that have duplicated or triplicated in Fol4287 are 1/2 and 1/3 lower respectively than in non-duplicated Fol4287 regions (indicated with green lines).

**Fig 2.**
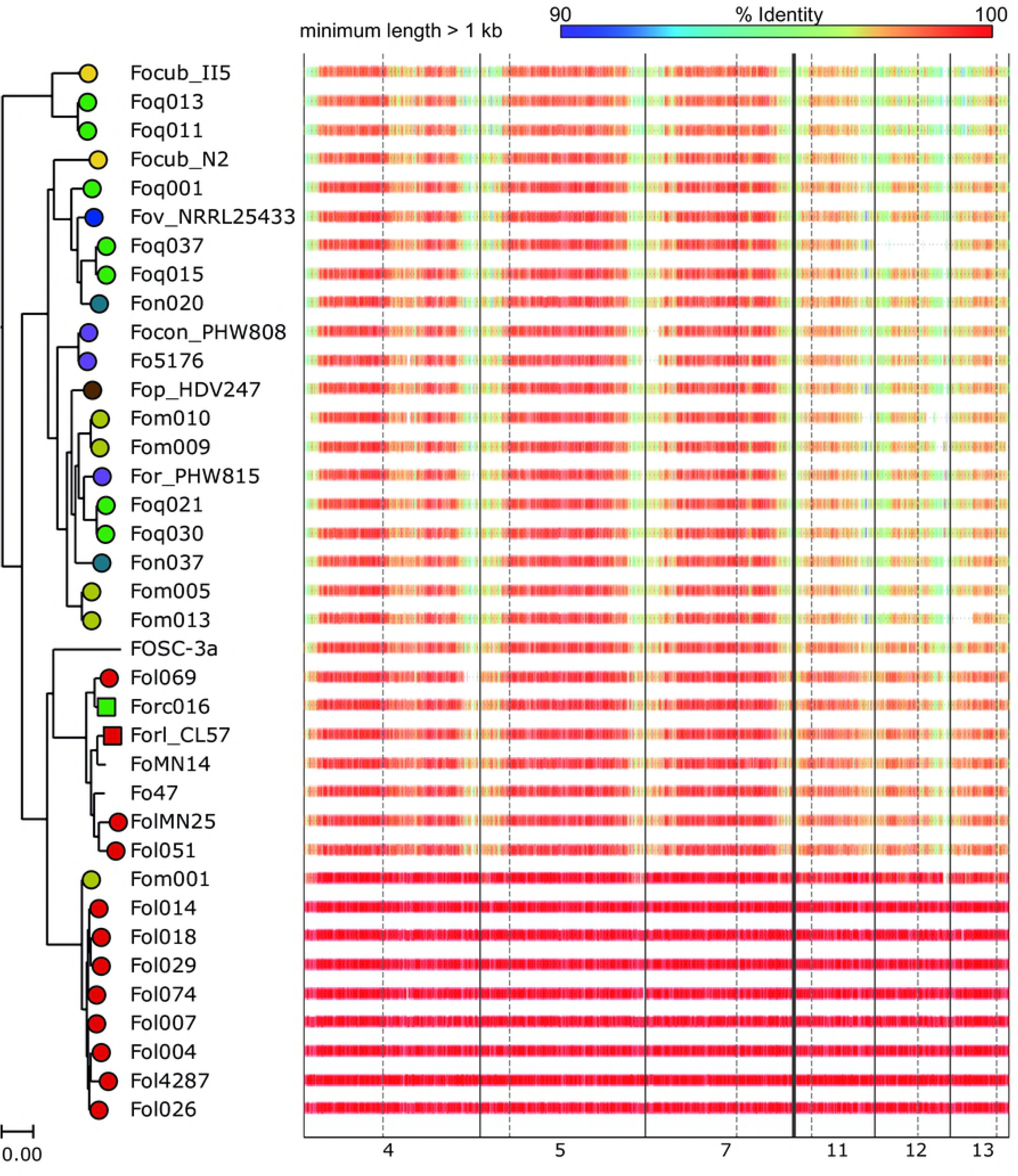
Core and fast-core chromosomes in the Fol4287 genome. Horizontal bars -colored according to the % identity in the alignment- indicate presence of Fol4287 core chromosomes 4,5,7,8,11,12, and 13 in 23 other *Fusarium oxysporum* isolates. Leaf nodes in the phylogenetic tree are colored according to *forma specialis*, isolates causing wilting symptoms are represented with a circle, those that cause root rot are represented with squares and those that are nonpathogenic do not have a shape or color. Only alignments that span more than 1 kb and are more than 90% similar are included. Within the Fol4287 clonal line, all chromosomes are mostly 100% identical, while when compared to other, more distant FOSC isolates, the three smallest core chromosomes, 11, 12 and 13, are more divergent than other core chromosomes (see also Figure S1). We denote these three chromosomes as fast-core chromosomes.

In conclusion, accessory chromosomes of Fol4287 largely correlate with phylogenetic clade, but chromosome 14 and regions on chromosome 3 and 6 correlate with host and are more conserved in sequence than core chromosomes.

We then asked whether these pathogenicity regions on chromosome 3 and 6 correspond to the small chromosome that was co-transferred with chromosome 14 in in the experiment described in [41], resulting in strains that were more virulent than the ones that received only one chromosome. We sequenced one strain that received chromosome 14 (from Fol007) and one - more virulent - strain that received chromosome 14 and the additional small chromosome. We mapped the reads thus obtained to the genome of Fol4287 and found that exactly those regions in Fol4287 that show hallmarks of horizontal transfer in our genome comparison (Fig 1A) correspond to chromosome 14 and the second, smaller ‘pathogenicity’ chromosome in Fol007 (Fig 1B).

Our results strongly suggest that horizontal transfer of chromosomes has indeed played an important role in the emergence of new pathogenic clonal lines in natural settings. Moreover, not all accessory regions in Fol4287 are alike: only some regions are associated with pathogenicity towards tomato. This suggests a division between accessory regions in Fol4287: chromosome 14 and part of chromosome 3/6 are involved in virulence towards tomato whereas the other accessory regions are not. From here on, we refer to the chromosomal regions associated with pathogenicity (i.e. chromosome 14 and part of chromosome 3 and 6) as ‘pathogenicity regions’, and to the rest of the accessory genome of Fol4287 as ‘lineage-specific regions’, as they only occur in strains that belong to the same lineage (vegetative compatibility group) as Fol4287.

### The core genome can be divided into regions with low divergence and regions with high divergence

To compare differences in dispensability among core chromosomes, we applied the same approach to the core genome of Fol4287. We found that all core chromosomes are present in all strains in our dataset, with the notable exception of chromosome 12 that is absent in the cucumber-infecting strain Foc037. This strain was included in several bioassays in a previous study, in which its virulence was confirmed [40]. Recently, flow cytometry was used to screen for loss of chromosome 1, 12 and 14 in Fol4287 [43]. In all three reported cases in which chromosome 12 was lost, no differences were observed in terms of virulence. Chromosome 12 is thus dispensable for growth as well as virulence, and should thus no longer be considered a core chromosome *sensu strictu*. In addition to complete loss of chromosome 12 in Foc037, we observed large deletions in core chromosomes in two melon-infecting strains: ∼0.5 Mb in a subtelomeric region of chromosome 12 in Fom010, and ∼0.5 Mb in a subtelomeric region of chromosome 13 in Fom013 (Fig 2).

In addition to differences in the propensity for loss or large deletions, there are striking differences in the level of sequence conservation within and among core chromosomes. The three smallest core chromosomes of Fol4287 – Chr. 11, 12 and 13 – are clearly more divergent than the other, larger core chromosomes (Fig 2) in terms of sequence similarity as well as synteny (Fig S2, Fig S3). Subtelomeric regions of all chromosomes and a ∼1 Mb central region of chromosome 4, associated with a genomic rearrangement when compared to *F. verticillioides* [41], also show elevated levels of sequence divergence and lower synteny levels (Fig S2, Fig S3). Notably, these chromosomes are not enriched in repetitive elements and have a similar gene density as other core chromosomes [41] (Fig S4). From here on, we will refer to chromosomes 11, 12 and 13 as ‘fast-core’ chromosomes.

### Genes on fast-core and accessory chromosomes have lower expression levels

We hypothesized that genes in different genome compartments exhibit differences in overall expression levels. To assess this, we queried an RNA-seq dataset that was generated to compare gene expression of Fol4287 grown in liquid culture (*in vitro*) to Fol4287 infecting tomato plants (*in planta*, 9 days post inoculation) [65] (Table S2). On the accessory genome, most genes -64% and 68% of genes on the lineage-specific and pathogenicity chromosomal regions respectively- are not expressed *in vitro*, (defined here as RPKM <= 0.1). On the fast-core and the core chromosomes, this is 46% and 25% respectively. *In planta*, 53% of genes on the accessory are not expressed (lineage-specific 51%, pathogenicity 57%), compared to <33% of genes on the fast-core and <19% of genes on the core chromosomes. Of the fast-core chromosomes, chromosome 12 - that is absent from Foc037 - has most genes that are not expressed in the conditions we tested (53% *in vitro*, 37% *in planta*). Overall, genes on the accessory genome and genes located in fast-core chromosomal regions (including the central region on chromosome 4) have lower expression than genes on the core genome, both *in vitro* and *in planta* (average over three biological replicates, Fig S5).

### Of the accessory genome, only chromosome 14 is enriched for genes that are differentially regulated during infection

We expected that the pathogenicity regions, required for virulence on tomato (Fig 1), would be enriched for genes that are upregulated during infection. When we compared *in vitro* to *in planta* expression levels, we found that chromosome 14 is indeed significantly enriched for genes that are upregulated during infection (P-value < 9.19 × 10^−10^ after Bonferroni correction, Fig 3, Fig S6). However, the ‘pathogenicity-related’ regions on chromosome 3 and 6 – although they correspond to the small chromosome in Fol007 that enhances virulence upon horizontal transfer – are not significantly enriched for upregulated genes. Like the other (lineage-specific) accessory regions, here only 3.1% of the genes were upregulated *in planta*, compared to > 14.7% of genes on chromosome 14. Hence of the pathogenicity regions, only chromosome 14 is enriched in genes that are differentially expressed during infection.

**Fig 3.**
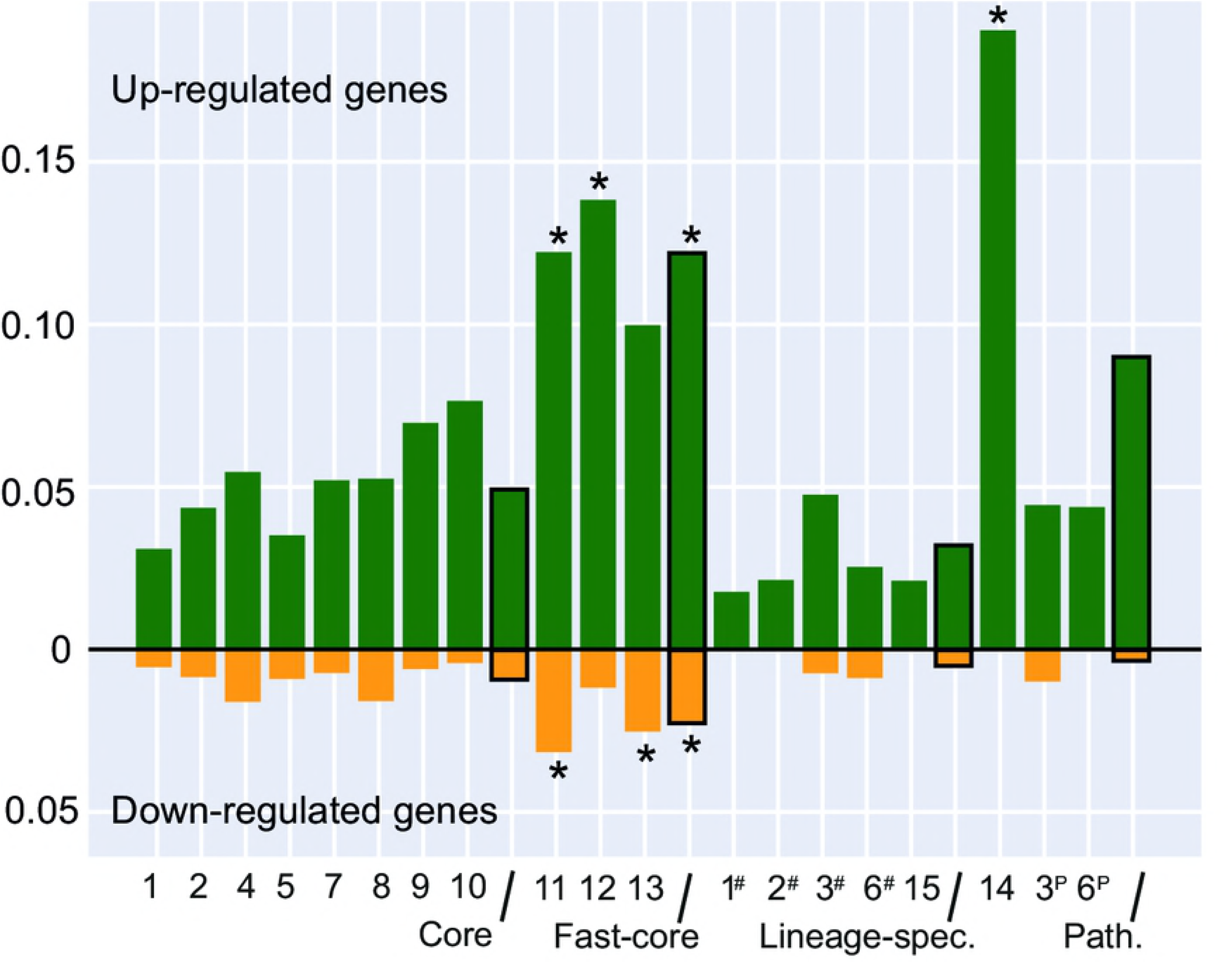
Fraction of up- and downregulated genes per chromosome. The bars represent the fraction of genes that is upregulated (green) or downregulated (red) *in planta* compared to *in vitro* per chromosome and per chromosome category (boxes with black lines). Significant enrichment (P-value < 0.001) of up- or downregulated genes with respect to the rest of the genome is indicated with asterisk signs. Lineage-specific parts of core (Chr. 1 and 2) or accessory (Chr. 3 and 6) chromosomes that are lineage-specific are denoted with a # on the x-axis.

We then asked whether the upregulated genes on chromosome 14 have similar functions as those encoded on chromosome 3 and 6. Of the 61 genes on chromosome 14 that are upregulated during infection, 16 encode small, secreted proteins, 13 of those are SIX (Secreted In Xylem) genes. About a third (19 out of 61) encodes enzymes, of which four are secreted. The remainder of upregulated genes includes three genes encoding for transcription factors, two of which are homologs of *FTF1*, that induces expression of effector genes [65], three putative transposable elements, a transporter, one encoding a membrane protein and one encoding a putative argonaute-like silencing protein (Table S3). The remaining 19 upregulated genes have no known functional domains and contain no secretion signal peptide. The proteins encoded by the 28 upregulated genes located in the pathogenicity regions on chromosome 3 and 6 include ten transposase-like proteins with a domain of unknown function, five enzymes and two small secreted proteins, one of which is a homolog of *SIX8* (Table S4). The three downregulated genes in the pathogenicity regions on chromosome 3 and 6 encode proteins with no known function and without a secretion signal peptide. In conclusion, more than a quarter of the upregulated genes on chromosome 14 encode small, secreted proteins, compared to less than 8 % of the upregulated genes on the pathogenicity regions on chromosome 3 and 6.

### Fast-core regions are enriched for genes upregulated during infection and are involved in metabolism, transport and defence

Interestingly, while on the ‘normal’ core chromosomes only 4% of the genes is differentially regulated during infection, on the fast-core chromosomes this is > 10%. Hence not only chromosome 14, but also the fast-core chromosomes are significantly enriched for differentially expressed genes (P-value < 1.17 × 10^−31^), both up- (P-value < 6.5 × 10^−25^) and downregulated (P-value < 1.67 × 10^−06^) (Fig 3). Not just fast-core chromosomes 11, 12 and 13, but also the other regions that showed elevated levels of sequence divergence (Fig 1, Fig 2, Fig S2) such as the subtelomeric regions of core chromosomes and the central region on chromosome 4 have more differentially expressed genes than core regions (Fig S6).

Compared to chromosome 14, the fast-core chromosomes are relatively conserved within the FOSC (Fig 1, Fig 2, Fig S2), suggesting that the upregulated genes on the fast-core chromosomes are involved in common infection-related processes while chromosome 14 contains host-specific genes. Of the 277 upregulated fast-core genes, 93 (∼34%) encode proteins that contain a putative signal peptide for secretion. This is similar to the proportion of upregulated genes on chromosome 14 that encode proteins with a predicted secretion signal peptide, but on chromosome 14 these are mostly small effector genes (Table S4). When we compared the functional annotations of upregulated fast-core genes to that of the rest of the genome, we find that upregulated fast-core genes are enriched for genes involved in metabolism, cellular transport, transport facilitation and transport routes and cell rescue, defense and virulence. More specifically, upregulated fast-core genes are enriched for genes involved in polysaccharide and other C-compound and carbohydrate metabolism, sugar, glucoside, polyol and carboxylate catabolism and extracellular polysaccharide degradation, in transport facilities, C-compound and carbohydrate transport and cellular import, and in degradation/modification of foreign (exogenous) polysaccharides (Table S5). The fact that these genes are differentially expressed during infection suggests that genes in fast-core regions are involved in digestion of plant material and import of products of digestion.

Not only fast-core genes that are upregulated during infection, are involved in metabolism, cellular transport, transport facilitation and transport routes and cell rescue, defense and virulence: when we also include fast-core genes that are not differentially regulated during infection, or are downregulated during infection, we find that these are involved in the same three broad functional categories (Fig S7) and in ‘Interaction with the environment’. Metabolic genes on the fast-core are typically involved in ‘metabolism of carbon compounds and carbohydrates’ (49.6% of metabolic genes) and ‘secondary metabolism’ (38.7% of metabolic genes). Transport genes are mostly involved in ‘transport of carbon compounds and carbohydrates’ (50.9% of transport genes), ‘cellular import’ (61.7% of transport genes) – in particular ‘non-vesicular cellular import’ (46.1% of transport genes) and ‘transport facilities’ (85.2% of transport genes). Defence genes are mostly genes involved in ‘detoxification’ (67.8% of defence genes) including ‘detoxification involving cytochrome P450’ (15.3% of defence genes) and in ‘disease, virulence and defence’ (32.2% of defence genes). This indicates that the clustering of genes that are involved in metabolism, transport and defence on fast-core chromosomes is not limited to genes that are involved in infection.

### Like accessory chromosomes, fast-core chromosomes are enriched with H3K27me3

Recent studies have revealed differences in chromatin-mediated regulation of core versus accessory chromosomal regions in pathogenic filamentous fungi: under *in vitro* conditions accessory chromosomes or chromosomal regions are enriched in H3K27me3, which is correlated with gene silencing, whereas core chromosomes are enriched in H3K4me2, which is correlated with gene activation [31,32,34]. To test whether the same holds true for *Fusarium oxysporum*, we performed ChIP-seq experiments for these two different histone marks (Table S6). We found that in *F. oxysporum*, similar to what was reported for *F. graminearum*, *F. fujikuroi* and *Z. tritici* [31,32,34,66], accessory regions are enriched for H3K27me3 and depleted in H3K4me2 (Fig 3, Fig S8). Interestingly, the same holds true for the fast-core chromosomes and sub-telomeric regions, in sharp contrast to the rest of the core genome that is enriched in H3K4me2 and depleted in H3K27me3 (Fig 4, Fig S8, Fig S9, Table S7).

**Fig 4.**
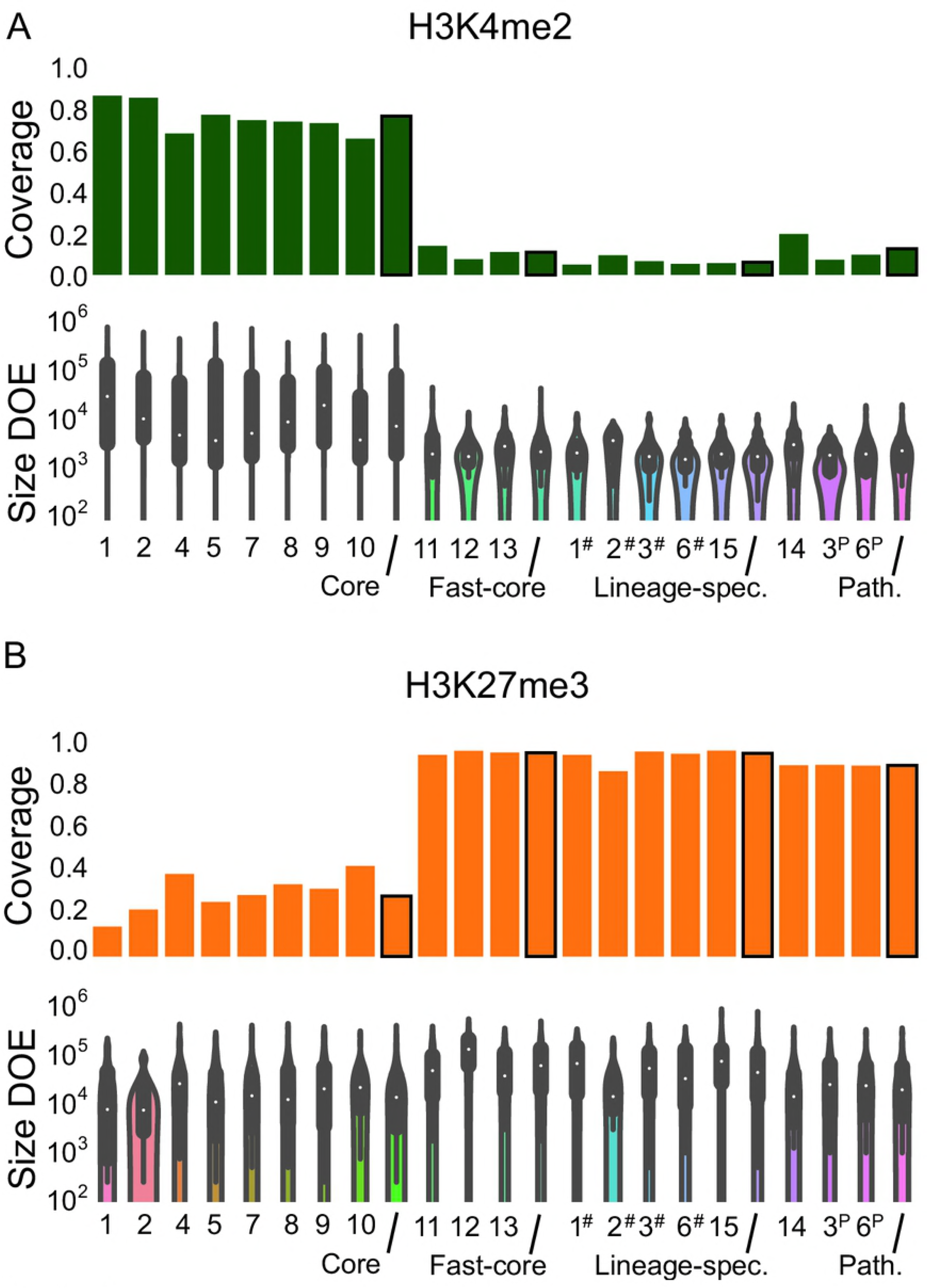
Core chromosomes are enriched for H3K4me2 and fast-core and accessory chromosomes for H3K27me3. We mapped ChIP-seq reads to the genome of Fol4287 and identified domains of enrichment (DOE). For each chromosome or chromosomal region (x-axis) we plot how much is covered in a DOE (top panels) and the size-distribution of DOEs (bottom panels, violin plots). Lineage-specific parts of core (Chr. 1 and 2) or accessory (Chr. 3 and 6) chromosomes that are lineage-specific are denoted with a ^#^ on the x-axis. The regions on chromosome 3 and on chromosome 6 that are associated with pathogenicity on tomato are denoted with a ^P^. Bars with thick lines represent the coverage per chromosome category (core, fast-core, lineage specific or pathogenicity) rather than per chromosome. A. Coverage and size distribution of DOE of H3K4me2. For most core chromosomes more than 50% is enriched for H3K4me3, while for the fast-core and accessory chromosomes, this is less than 20%. The DOE of H3K2me2 are larger on the core chromosomes than in the other regions. B. Coverage and size distribution of DOE of H3K27me3. For most core chromosomes less than 50% is enriched for H3K37me3, while for the fast-core chromosomes more than 80% is covered in H3K27me3 and the lineage-specific and accessory chromosomes are more enriched in H3K27me3 than the core chromosomes. In general, the size of the H3K27me3 DOEs is larger than that of H3K4me2 DOEs (note that y-axis is in log-scale).

H3K4me2 was found in or close to coding regions (Kendall’s Tau <= 0.39, *p* < 1 × 10^−300^), and is negatively correlated with the presence of transposons (Kendall’s Tau <= −0.33, *p* < 1 × 10^−300^). On the core genome, most genes and promoter regions overlap with an H3K4me2-enriched domain, while most transposons overlap with an H3K27me3-enriched domain (Fig S10). H3K4me2-enriched domains on core chromosomes are larger than those on fast-core and accessory chromosomes: they span between 10 and 100 kb on the core genome, compared to between 2.5 and 5 kb on the fast-core and accessory chromosomes (Fig 4, Fig S9, Table S7). More than 49% of the core genome is covered in H3K4me2-enriched domains, while only 2.5%-13% of the non-core genome is part of an H3K4me2-enriched domain.

In contrast, more than 72% of the accessory genome, ∼93% of the fast-core genome and <27% of the core genome is part of an H3K27me3-enriched domain. H3K27me3 is negatively correlated with the presence of coding sequences (Kendall’s Tau <= − 0.25, P < 3 × 10^−180^) and is enriched near or in transposons (Kendall’s Tau >= 0.35, P < 5,9 × 10^−283^). In general, H3K27me3 occurs in larger blocks than H3K4me2, on all chromosomes (Fig 4, Fig S9, Table S7). The average size of an enriched domain ranges between 15 to 60 kb on core chromosomes, where on the fast-core chromosomes domains vary between 57 and 146 kb (domain size averaged per chromosome). Especially on chromosome 12, H3K27me3-enriched domains are large: averaging ∼146 kb in one experiment and ∼83 kb in the other (Fig 4, Fig S9, Table S7).

H3K27me3 is associated with facultative heterochromatin, hence we expect low expression of genes that reside in a region that is enriched in this histone modification. In contrast, H3K4me2 is associated with euchromatin, hence we expect high expression levels for genes that are located in regions enriched in this histone mark compared to genes that reside in H3K37me3 domains of enrichment. As mentioned above, the fast-core and accessory chromosomes -enriched in H3K27me3- have more genes with low expression *in vitro* and -to a lesser extent- *in planta* than core chromosomes. Genes that are located in regions that are associated with H3K27me3 have lower transcript levels, both in *in vitro* and in *in planta* conditions (P value ∼ 0, Welch’s t-test on ranked data), while genes that are located in regions associated with H3K4me2 have higher transcript levels than the rest of the genome (P value ∼ 0, Welch’s t-test on ranked data), consistent with the assumed role of these histone marks in gene silencing and activation in other fungi [31-34,66]. When we compared gene expression *in vitro* to *in planta* conditions, we found that most transcriptional changes involve *up*-*regulation* of genes: 896 out of 1036 differentially expressed genes are upregulated *in planta* compared to *in vitro* (>86%) (Table S4, Fig S6). Of these upregulated genes, more than 66% is located in an H3K27me3-enriched domain (P-value < 1.6 × 10^−41^, hypergeometric test). The differences in histone codes of core, fast-core and accessory genomes may allow for large-scale coordinated expression of genes that are involved in life-style switches, such as infection.

### Fast-core chromosomes are not enriched for genes under positive selection

The differences in functional gene classes enriched in core, fast-core, accessory and pathogenicity chromosomes raises the question to what extent the observed hierarchy in sequence divergence levels (Fig 1, Fig 2) of these four categories of chromosomes can be explained by positive selection. A previous study identified genes under positive selection based on multiple sequence alignments of genes that belong to nine Fusarium species, including *F. oxysporum*, comparing synonymous and non-synonymous substitution rates and testing for the occurrence of site-specific balancing selection. This study revealed that all Fol4287 accessory chromosomes (pathogenicity and lineage-specific regions) are enriched for genes that are under positive selection [24]. The core and fast-core chromosomes contain relatively few genes that are inferred to be under positive selection: 308 (< 2.7% of tested core genes) and 66 (< 2.8% of tested fast-core genes) respectively (Table S8), compared to 192 genes located in accessory regions (> 9.8% of tested accessory genes; 8.4% lineage-specific and 12.5% pathogenicity). Differentially expressed genes are not significantly enriched for genes under positive selection (P-value > 0.5). Hence positive or diversifying selection on genes that are located on the fast-core chromosomes does not explain the high levels of sequence divergence we observed.

We then hypothesized that genes on these chromosomes are under relaxed negative selection, which could also explain differences in sequence divergence: many genes have low expression levels. Moreover, fast-core chromosome 12 can be lost in both Fol4287 and Foq037 without losing virulence [40,43] (Fig 2). Per gene, we inferred the number of synonymous substitutions per synonymous site (d_S_), non-synonymous substitutions per non-synonymous site (d_N_), as well as their ratio (d_N_/d_S_) based on alignment with their bidirectional best BLAST hit in *F. verticillioides*. We found that higher substitution rates in genes located on fast-core chromosomes compared to those located on other core chromosomes (P-value < 2.8e-39, P-value < 8e-307, P-value < 0.0015, for d_S_, d_N_ and d_N_/d_S_ respectively, Welch’s t-test on ranked data, Fig 5A, Fig S11). Synonymous substitutions should be little influenced by relaxed or positive selection, yet we observed also a significantly higher *synonymous* substitution rate on fast-core chromosomes. We therefore conclude that relaxed selection on dispensable genes and genes with low expression levels alone, does not explain differences in sequence divergence between fast-core chromosomes and other core chromosomes.

**Fig 5.**
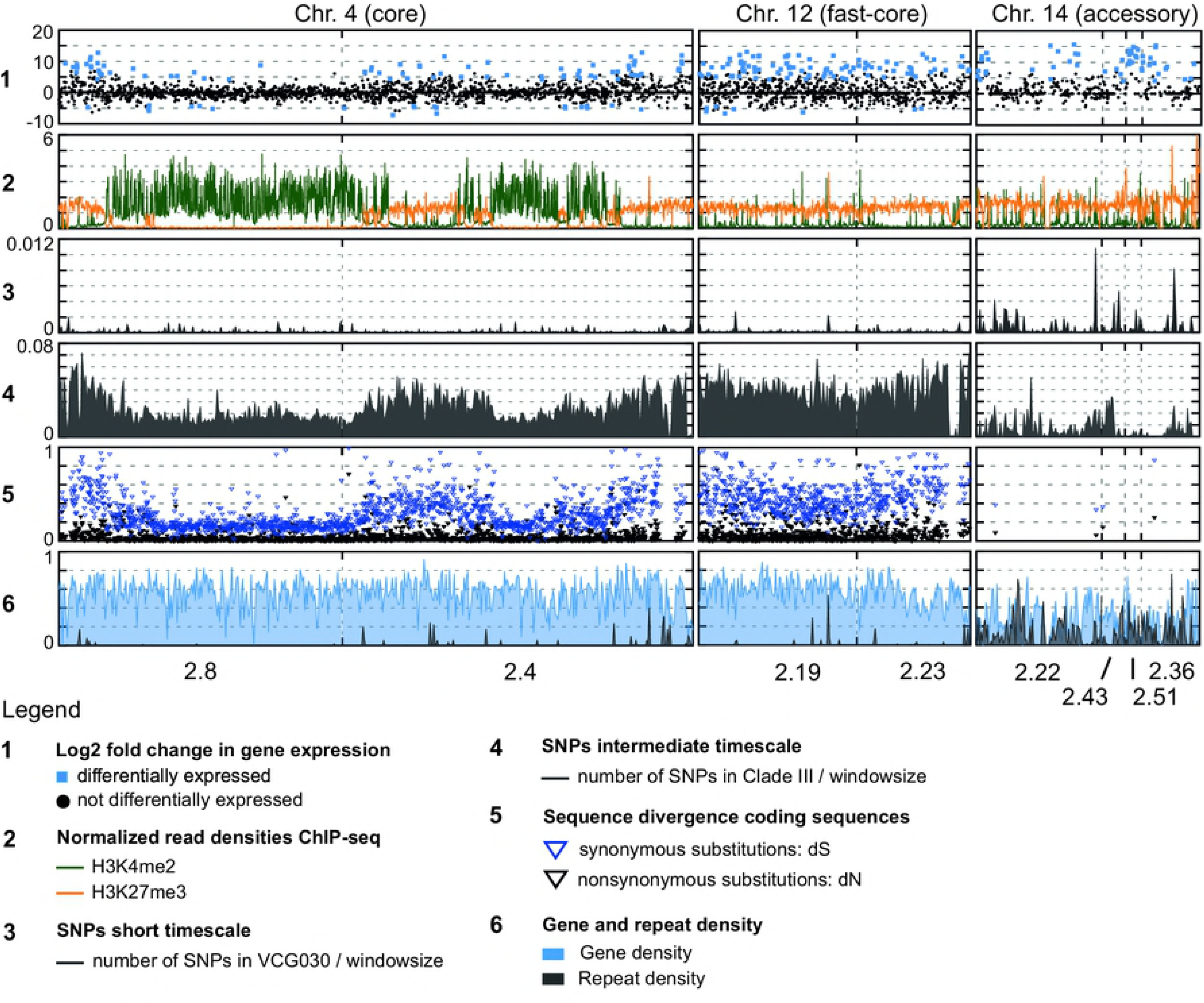
H3K27me3-enriched regions have higher levels of sequence divergence, independent of repeat density. These three examples, one core, one fast-core and one accessory chromosome, demonstrate how presence of H3k27me3 (panel 2) coincides with differential expression (panel 1), elevated levels of sequence divergence (panel 3-5), but not necessarily with differences in either gene- or repeat density. The genes on core chromosome 4 that are differentially expressed during infection (panel 1, blue squares) are located mostly in the sub-telomeric regions and one central region. These regions are enriched in H3K27me3 (panel two, orange) and depleted in H3K4me3 (panel two, green). The fast-core chromosome has many differentially expressed genes, most of which are upregulated during infection. There are no large regions that are either enriched or depleted in differentially expressed genes. The same holds true for H3K27me3, which is relatively equally distributed along the chromosome. More or less the same holds true for chromosome 14, the pathogenicity chromosome and the only accessory chromosome that is enriched for genes that are upregulated during infection. Here also H3K27me3 is relatively uniformly distributed along the chromosome: unlike core chromosomes, H3K27me3-ated regions are not interrupted by regions with high levels of H3K4me2. When looking at sequence divergence on a very short timescale (panel 3, density of SNPs called in strains that belong to VCG030) there are no distinct differences within and between the core and fast-core chromosomes, while chromosome 14 clearly has a higher SNP density (see Figure S14 for more detail). In contrast, on longer timescales (panel 4, density of SNPs called in strains that belong to clade III, panel 5, d_N_ (black) and d_S_ (blue) based on bidirectional best hits with sister species *F. verticillioides*), we can clearly see that regions that are enriched in H3K27me3 on the core chromosome correspond with regions with higher d_N_ and d_S_ (see also Figure S14 and Figure S15) and a higher SNP density. Fast-core chromosomes resemble Sub-telomeric core regions. Note that gene- and repeat density are relatively uniform on the core and fast-core chromosomes (see also Figure S4). The SNP density of accessory chromosomes cannot directly be compared to the core and fast-core chromosomes because they are lost in many strains in Clade III and in *F. verticillioides*. The accessory chromosomes are depleted in genes and enriched in repeats.

### Rate of sequence divergence is elevated in H3K27me3-enriched regions on core chromosomes

Interestingly, both genes on fast-core chromosomes and genes on core chromosomes that are located close to a telomere also have higher d_S_ and d_N_ values (Fig 5A, Table S8, Fig S11). When we compare substitution levels for genes that are located in a region enriched in H3K27me3 on core chromosomes, the distribution of d_N_ and d_S_ of these genes is similar to those located on the fast-core chromosomes (Fig 5A, Fig S12). If we view the synonymous substitution rate as an approximation of the molecular clock, these results suggest that this “clock is ticking faster” in regions that are enriched in H3K27me3.

To study differences in rates of sequence evolution in more detail we inferred Single Nucleotide Polymorphisms (SNPs) on two timescales. The cumulative SNP density of the ten tomato-infecting strains that belong to the same VCG as Fol4287 (the bottom tomato-infecting lineage in Figs 1 and 2, see Materials and Methods) approximates the accumulation of mutations on a short timescale. Assuming no horizontal transfers have occurred within this lineage, we can compare accessory chromosomes with core and fast-core chromosomes, as all strains possess Fol4287’s accessory chromosomes, albeit not all complete chromosomes. We found that the accessory chromosomes accumulated more SNPs than the fast-core and core chromosomes, but that the SNPs density of core and fast-core chromosomes is similar (P-value > 0.35, hypergeometric test, Fig 5, Fig S13, Fig S14). We found that on fast-core chromosomes SNP density was higher in H3K27me3-enriched regions (comprising more than 90% of the fast-core chromosomes), compared to regions that are not enriched in H3K27me3, but that this difference was not significant. However, on core chromosomes H3K27me3-enriched regions have a significantly higher density of SNPs (P-value < 1.5 × 10^−7^). When we included SNPs in less closely related strains (i.e. all strains in the Clade III, the clade depicted in Fig 1), we found that fast-core chromosomes, the central region on chromosome 4 and sub-telomeric core regions have more SNPs than (other) core regions (Fig S13). Hence, on a longer timescale, as observed when comparing genome sequences, we find that regions that are enriched in H3K27me3, have higher sequence divergence (P-value ∼ 0, hypergeometric test, for both core and fast-core chromosomes). Due to the fact that the accessory chromosomes are absent from a number of strains in Clade III, and the fact that the pathogenicity chromosomes were obtained through horizontal transfer, we cannot directly compare accessory chromosomes to core and fast-core chromosomes. We conclude that, especially on core chromosomes and/or on longer timescales, H3K27me3-enriched regions accumulated more SNPs compared to regions that are not enriched in H3K27me3.

We calculated correlations between change in gene expression, sequence divergence and histone modifications on core or fast-core chromosomes, and found two clusters -each corresponding to a different ‘speed’- where repeat density and SNP density cluster with H3K27me3, while gene density and gene expression levels cluster with H3K4me2 (Fig 6A and B). Interestingly, H3K27me3 correlated most strongly with synonymous substitution rates and SNP densities in Clade III, and much less with repeat density. For accessory chromosomes this distinction was not so clear; we observed mostly very low correlation coefficients between different properties (Fig 6C). In conclusion, fast-core and core chromosomes harbour similar “speeds”, but in different quantities. The evolutionary processes underlying the different “speeds” on core and fast-core chromosomes probably have had less influence in shaping the organization of accessory chromosomes.

**Figure 6.**
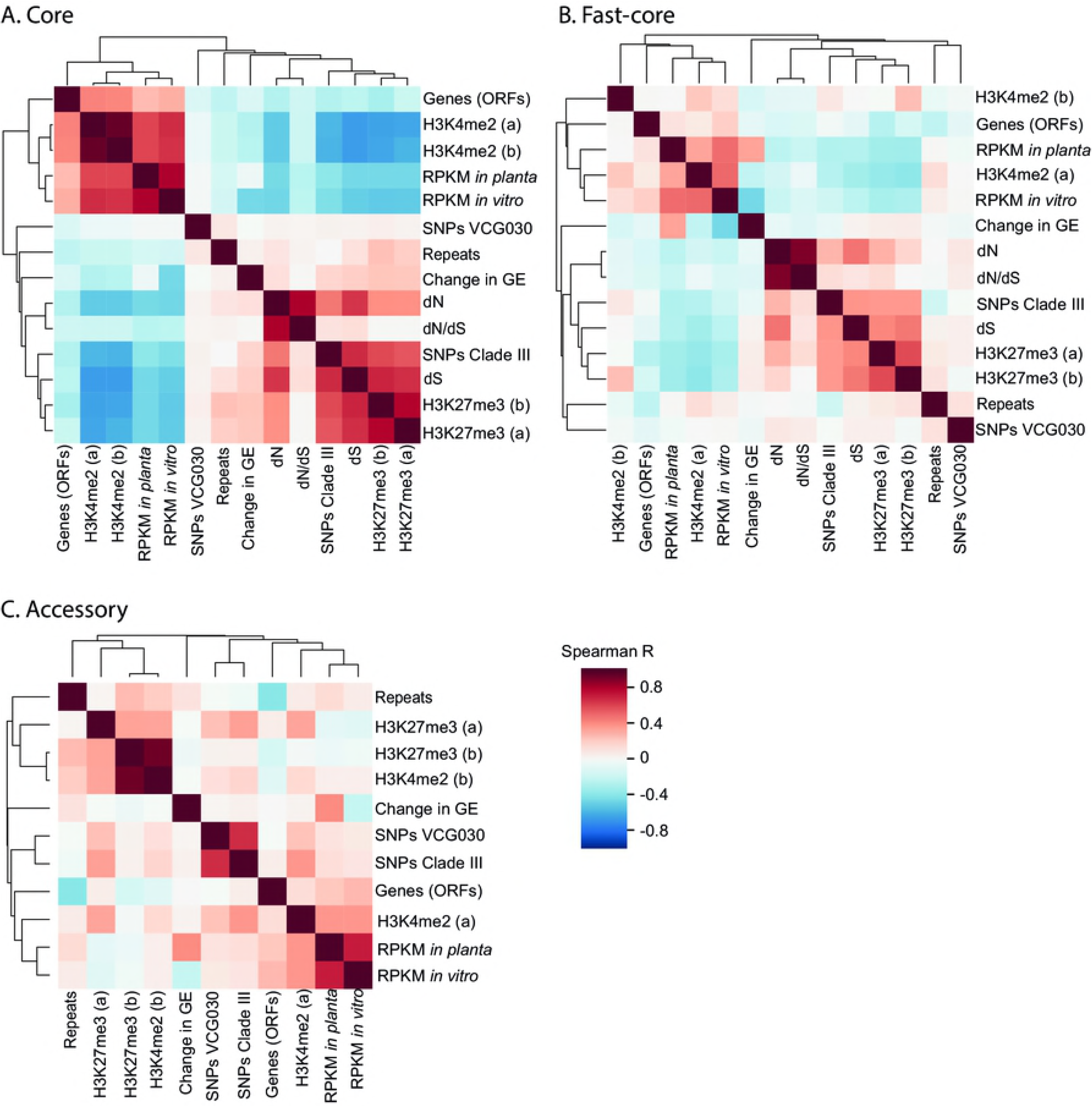
Clustering of genome characteristics. We calculated the following genome characteristics for 6186 10 kb non-overlapping windows across the entire Fol4287 genome: **Genes (ORFs**): the number of bp that are part of a coding region divided by the window size, **Repeats**: the number of bp that are part of a repeat region divided by the window size, **RPKM *in planta***: mean (over 3 replicates) RNA-seq read counts for the *in planta* experiment, **RPKM *in vitro***: mean (over 3 replicates) RNA-seq read counts for the *in vitro* experiment, **Change in GE**: log2 fold change in gene expression *in planta*/*in vitro*, **H3K4me2 (a) and (b)**: average read density for the two replicates of the H3K4me2 ChIP-seq experiment, **H3K27me3 (a) and (b)**: average read density for the two replicates of the H3K27me3 ChIP-seq experiment, **dN**: average number of nonsynonymous substitutions per nonsynonymous site dN, **dS**: the average number of synonymous substitutions per synonymous site dS, **dN/dS**: average ratio between synonymous and nonsynonymous substitutions, **SNPs VCG030**: density of SNPs in VCG030 with respect to Fol4287, **SNPs Clade III**: density of SNPs in clade III with respect to Fol4287. Spearman correlation coefficients (R) are depicted in a heatmap, genome characteristics were clustered with average linkage using 1-R as a distance measure. A. Clustering based on core chromosomes. The clustering shows two regimes: one gene-rich, highly expressed, H3K4me2-ated and one repeat-rich, H3K27me3-ated with high levels of sequence divergence. B. Clustering based on fast-core chromosomes (11, 12, 13). As with core chromosomes, the clustering shows two regimes: one gene-rich, highly expressed, H3K4me2-ated and one repeat-rich, H3K27me3-ated with high levels of sequence divergence, albeit less pronounced as correlation coefficient tend to be lower. C. Clustering based on accessory chromosomes. Unlike the core and fast-core chromosomes, levels of sequence divergence correlate with gene density and expression level. However, this may be an artefact from the SNP calling process, in which reads that map to multiple locations -e.g. reads that map to transposons- are excluded. Due to lack of data points, dN, d_S_ and dN/dS were excluded.

## Discussion

In this study, we found that *F. oxysporum* has a more complex genome organization than previously proposed [41]. While it has been known for several years that *F. oxysporum* has core and accessory chromosomes, our more detailed comparisons revealed distinct subcategories. Core chromosomes can be divided into ‘core’ (*sensu strictu*) that are indispensable, conserved, enriched in H3K4me2 and not enriched in genes that are differentially expressed during infection and ‘fast-core’ that are conditionally dispensable, less conserved, enriched in H3K27me3 and enriched in genes that are differentially expressed during infection. Similarly, we suggest to classify the accessory genome, which is enriched in H3K27me3, into lineage-specific chromosomes that are specific to a clonal line and pathogenicity chromosomes that are present in all strains that infect a certain host. Apart from distinct chromosomes, we also observed regions -parts of chromosomes- that could be assigned to a different category as the rest of the chromosome, such as sub-telomeres, or translocated segments on *bona fide* core chromosomes that can be classified as fast-core, i.e., segments of core chromosomes 1 and 2 that are lineage-specific, and segments of chromosome 3 and 6 that are considered pathogenicity regions.

The genome alignments and the analyses of SNP densities along the genome showed that the different subcategories can be ordered into a hierarchy with respect to their rate of sequence divergence: compared to core regions, fast-core regions accumulated more SNPs and alignments of fast-core regions with genome sequences of other FOSC isolates have lower percent identity. Fol4287’s accessory regions are absent from most strains and accumulated many SNPs on a timescale, on which we found no significant difference between core and fast-core chromosomes. Similarly, synteny is most conserved in core regions, less in fast-core regions, and even less in pathogenicity regions and in lineage-specific regions.

Interestingly, we find that -to some extent- different levels in this hierarchy correspond to levels in host-interactions: both fast-core chromosomes and chromosome 14 are enriched for differentially expressed genes, but genes on the less volatile fast-core chromosomes are typically involved in metabolism, transport and defence, whereas chromosome 14 contains many effector genes that are hypothesized to interact with host proteins [67]. The fast-core chromosomes are in general enriched in genes involved in metabolism, transport and defence, importantly, including genes that are not differentially regulated during infection. *F. oxysporum* is not an obligate pathogen and we predict that levels in the observed sequence divergence hierarchy also correspond to different levels on other functional ‘axes’, including interactions with other soil organisms, where for example lineage-specific regions may play a more important role.

Although we were able to associate different levels of sequence divergence with different levels of host interaction, this correspondence is not perfect. While pathogenicity chromosome 14 of Fol4287 is enriched in genes with induced expression during infection, the regions on chromosome 3 and 6 are not, neither do these regions encode any known effectors. Yet these regions are important for virulence, as combined horizontal transfer of this region with chromosome 14 increases the aggressiveness of the recipient strain compared to strains that receive only chromosome 14 [41]. Moreover, our results show that these regions are present in all tomato-infecting strains, although possibly as a single copy. A remaining question is thus whether these regions on chromosome 3 and 6 contains one or several genes that are important for infection and whether the rest of these regions is ‘hitchhiking’. In comparison, Fol strains that have lost large segments of chromosome 14, including *SIX6*, *SIX9* and *SIX11*, whose products have been found in the xylem sap of infected tomato plants [68], are not significantly less virulent than strains with a complete chromosome 14 in laboratory conditions [43]. This suggests that segments of chromosome 14 may be hitchhiking with genes that are important for infection and that reside on separate regions of chromosome 14, such as *SIX1*, *SIX3* and *SIX5* [69,70]. On the other hand, the genes that may seem redundant for virulence in conditions tested in bioassays may perform a role in other conditions. For example, *SIX6*, found in multiple FOSC strains as well as in *Fusarium hostae* and *Colletotrichum* species, is required for virulence of *F. oxysporum* f. sp. *radicis-cucumerinum* towards cucumber only at relatively high temperatures [64]. Despite the fact that genetic hitchhiking and incomplete functional characterizations obscure relations between function and sequence divergence, we were still able to associate physical - with functional clustering, as observed on fast-core regions on core chromosomes, on fast-core chromosomes and on chromosome 14.

We not only found that pathogenicity chromosomes are specific to tomato-infecting strains, we also found that horizontal chromosome transfer (HCT) most likely played an important role in distributing the pathogenicity regions among different FOSC strains. Population genetic studies in FOSC revealed that pathogenic strains group with non-pathogenic ones [45,48,49,52-54,71-73], consistent with a scenario in which a non-pathogenic strains receives at least part of a pathogenicity chromosome and becomes pathogenic on a specific host. This is more likely to occur than switching from one host to another because first of all, non-pathogens are more abundant and secondly, pathogens may secrete effectors that enhance infection in one host but are recognized in another. The importance of HCT in the emergence of new disease depends, among other factors, on how long a pathogenicity chromosome can remain intact in a population. The small pathogenicity chromosome in Fol007 is part of chromosome 3 and chromosome 6 in Fol4287. Interestingly, no horizontal transfer of chromosome 3 and 6 has been observed in the lab [63]. Although we found that chromosome 14 and part of chromosomes 3 and 6 are present in all tomato-infecting strains included in our dataset, whether these segments are present as a single chromosome, multiple chromosomes or regions attached to other chromosomes, remains to be investigated. The presence of transposons impedes genome assembly from short reads, making it difficult to resolve this question with currently available genome sequences. However, we can conclude that the tomato-infecting strains in our dataset probably arose from a non-pathogenic strain after obtaining one or multiple pathogenicity chromosomes. We found no evidence for horizontal transfer of fast-core chromosomes in this comparison. Nevertheless, given the fact that genes on fast-core chromosomes are involved in infection and that we have observed transfer of core chromosomes in high-throughput experiments [63], we predict that horizontal transfer of core and fast-core chromosomes can also occur in natural settings. Co-transfer of fast-core chromosomes with accessory chromosomes may in some cases even enhance pathogenicity of the recipient strain.

HCT, or -an extreme case- hybridization, can have an enormous evolutionary impact. As we discussed above, it allows for fast spreading of fitness-enhancing pathogenicity chromosomes. Moreover, it provides a means to escape “Muller’s ratchet” in mainly asexual species by exchanging at least parts of core chromosomes. The transposons that reside on pathogenicity chromosomes can play a role in fast adaptation to host-resistance, [19,74] probably mostly by mediating rearrangements [5,8,75], thus potentially disrupting modular genomic organizations and creating novel chromosome segments. Moreover, transposons may disperse into core chromosomes [76] where the chance of harmful effects of transposon insertion is larger than on dispensable chromosomes. This interplay between beneficial and detrimental effects of transposons and mobile and/or accessory chromosomes on their host genomes has not yet been systematically investigated, even though it may strongly influence evolutionary outcomes, including the emergence of novel pathogens by host jumping. Population genetic studies and experimental evolution, combined with long read sequencing, may shed light on these important issues. We demonstrated here how HCT events can be detected and visualized using whole genome sequencing.

We found that none of the core chromosomes of Fol4287 are rich in transposons (Fig S4), but the three smallest core chromosomes, termed ‘fast-core’, are distinct from the other core chromosomes in terms of sequence divergence, synteny, dispensability, and selected histone modifications. In that respect they are similar to for example accessory regions in *Fusarium graminearum*, that have a higher SNP density, are enriched in H3K27me3, yet are neither depleted in coding sequences nor enriched for repeats [30]. There has been much debate on the influence of chromatin states on *de novo* mutation rates. Constitutive heterochromatin may be less accessible to the DNA repair machinery. Research in cancer cell lines and mammalian germ line cells yielded conflicting results [77]. A recent study in *Schizosaccharomyces pombe* revealed that in this fungus the DNA mismatch repair machinery favours euchromatin. Similar to what we found here (Fig 5, Fig 6), the authors observed a higher mutation rate in (in their case H3K9me2-enriched) heterochromatin [78]. The fact that on core and fast-core chromosomes, H3K27me3 strongly associates with higher synonymous substitution rates (Fig 6A, B) and with density of SNPs accumulated on an intermediate timescale, suggests that H3K27me3 is correlated with or actively enhances mutation rates. However, this effect appears insufficient to generate noticeable differences in density of SNPs accumulated on very short timescales. Moreover, we do not find a strong correlation between H3K27me3 and SNP density on accessory chromosomes, which accumulated many more SNPs than fast-core chromosomes with comparable levels of H3K27me3. Accessory chromosomes, particularly pathogenicity chromosomes of strains that infect crops, may experience different selection pressures than core and fast-core chromosomes, which could affect sequence evolution. Additional histone modifications, possibly associated with a high density of repeats, may also further influence mutation rates.

Genome organization is shaped by a combination of intrinsic molecular mechanisms and natural selection. It is still not completely clear to what extent physical clustering of genes in the same functional category has a selective advantage, such as robust and stable co-regulation, efficient transferability of co-functional genes or increased evolvability due to locally increased mutation rates (including rearrangements), e.g. due to the presence of transposons or due to histone modifications that influence DNA repair efficiency. We here defined two subcategories of the core and the accessory genome and more fine-grained classifications are possible and probably valuable. Our subclassifications reveal a hierarchical modular genome organization - as is found with proteins and protein networks and is associated with evolvability [79-81]. A more detailed classification of genome ‘modules’, identifying more than two or four speeds, may help to disentangle the different molecular mechanisms, selection on the organism level -fitness- and long-term selection on genome structure - evolvability.

## Materials and methods

### Whole genome alignments

We used nucmer from the MUMmer package (version 3.23) to align the genome of Fol4287 to the 58 FOSC genomes [40] in our dataset (Table S1, (https://github.com/LikeFokkens/whole_genome_alignments). We flagged – maxmatch to allow for more than one alignment per region to be able to identify duplicate regions, other than that we used default settings. We used show-coords with default settings (no filtering) to obtain tab-separated files with aligned regions and custom Python scripts to plot the percent identity (Figs 1, 2, and S2) or length (Fig S3) of these aligned regions (https://github.com/LikeFokkens/genome-wide_plots).

### Illumina sequencing of horizontal chromosome transfer strains

Two strains, Fo47-1A that obtained one chromosome and Fo47-2A that obtained two chromosomes in the horizontal transfer experiment [41] were stored at −80 °C and revitalized on potato dextrose agar (PDA) plates at 25 °C. An overgrown agar piece was used to inoculate 100 ml NO_3_-medium (0.17% yeast nitrogen base, 3% sucrose, 100 nM KNO_3_). After 5 days at 25 °C shaking at 250 rpm, mycelium was harvested in miracloth (Merck, pore size of 22-27 μM) and dried overnight in the freeze dryer. Genomic DNA was isolated from freeze-dried mycelium as described in [36] and [40] and sequenced on a Illumina Hiseq2000 sequencer at the Beijing Genomics Institute.

### Read densities of DNA sequencing reads

We removed adapter sequences and low quality reads from reads obtained from Illumina sequencing as described above with fastq-mcf with ‘-q 20’ and a fasta file with all Illumina adapter sequences. We then first mapped these reads to the genome of Fo47 (the acceptor strain in the experiment), extracted the unmapped reads and mapped these unmapped reads to the genome of Fol4287 to identify the regions in Fol4287 that correspond to the chromosomes that were transferred from Fol007 (the donor strain in the experiment). For each strain (Fo47-1A and Fo47-2A), we mapped two libraries, one with insert size 170 bp, and one with insert size 500 bp. We used Bowtie 2 (version 2.2.6) [82], to map reads to the Fo47 genome, applying the following options: ‘‐‐end-to-end ‐‐fr –reorder’, for library with insert size 170 we added the options ‘-I 140 -X 200’ and for the library with insert size 500, we added the options ‘-I 400 -X 600’. We extracted the unmapped reads using samtools view (version 1.3.1), once with options ‘-Sbf 4 -F264’ to extract unmapped reads for which the mate was mapped to a samfile, once with options ‘-Sbf 8 -F260’ to extract mapped reads for which the mate was unmapped to a samfile, and once with options ‘-Sbf 12 -F256’ to extract unmapped reads for which the mate was also unmapped to a samfile. We sorted the samfiles using ‘samtools sort’ and merged them into a single using ‘samtools merge’. We used bedtools (version v2.24.0) ‘bam2fastq’ to convert the merged bamfile to fastq files. These fastq files were then mapped to the genome of Fol4287 using Bowtie2 with the same options as were used for mapping to the genome of Fo47. We removed putative PCR duplicates from the resulting bamfiles using Picard tools version 1.134 (http://broadinstitute.github.io/picard, MarkDuplicates) and used a custom Python script to calculate read densities for 10 kb windows and plot these on the genome (Fig 1B) (https://github.com/LikeFokkens/genome-wide_plots). Reads have been uploaded to ENA (accession PRJEB29294).

### Repeats

Repeats were identified using RepeatMasker (Repbase Libraries 20140131) with ‘–species ascomycota’ and otherwise default options. Repeats labeled as ‘Low_complexity’, Simple_repeat’, ‘rRNA’ or ‘Satellite/5S’ were filtered out. Repeat densities were calculated for 10 kb non-overlapping sliding windows using ‘bedtools coverage’.

### Functional enrichment analyses

We generated lists of (upregulated) genes located on fast-core chromosomes and in pathogenicity regions. We used the FungiFun tool [83] to predict functional enrichment and depletion (https://elbe.hki-jena.de/fungifun/fungifun.php), we chose ‘Fusarium oxysporum f. sp. lycopersici (strain 4287 / CBS 123668 / FGSC 9935 / NRRL34936)’ as species, and set adjusted (∼ after FDR correction) P-value < 0.001 as significance level (both under Advanced options). Accessory chromosomes typically contain genes with unknown function and less than 14% of genes in pathogenicity regions are annotated in FungiFun (Table S5C). Therefore we relied on the annotation published in [62] for chromosome 14. In addition we used HmmerWeb version 2.24.1 to predict Pfam domains (hmmscan, default settings) and the presence of secretion signal peptides. For the upregulated genes on the fast-core chromosomes we predicted secretion signal peptides via the SignalP 4.1 server, using default settings.

### Gene expression analyses

To compare *in vitro* and *in planta* gene expression levels, we used a dataset that has been described previously, hence we refer to [65] for an extensive description of methods. We used hypergeometric tests as implemented in scipy with Bonferroni correction to determine whether chromosomal regions are enriched for differentially expressed genes.

### ChIP-seq experiments and data analyses

*Fusarium oxysporum* f. sp. *lycopersici* 4287 (FGSC9935) was stored as a monoconidial culture at −80 °C and revitalized on potato dextrose agar (PDA) plates at 25 °C. An overgrown agar piece was used to inoculate 100 ml NO3-medium (0.17% yeast nitrogen base, 3% sucrose, 100 nM KNO_3_). After 3-5 days at 25 °C shaking at 250 rpm, spores were harvested by filtering through miracloth (Merck, pore size of 22-27 μM). Spores were centrifuged for 10 minutes at 2000 rpm, resuspended in sterile MilliQ water and counted [40]. We inoculated 100 ml of fresh NO3-medium with 10^7^ spores and cultured for 2 days at 25 °C and shaking at 250 rpm.

The ChIP-seq protocol [84] was similar to those used for *S. pombe* [85] and *Neurospora crassa* [86,87]. Here we only refer to important changes made for *F. oxysporum* strains. To the 2-day old cultures 20% formaldehyde (final concentration ∼0.8%) was added directly to the flask and incubated for 15 min at room temperature while shaking at 100 rpm. After formaldehyde quenching [84], 100-200 mg of the cell pellets were ground in liquid nitrogen. Too much cell mass will decrease ChIP efficiency. Digestion with micrococcal nuclease to release mostly mono- or dinucleosomes was as described [84] but should be optimized for different strains. ChIP-seq libraries were generated as described [84], with Illumina TruSeq adapters, libraries were size-selected between 200-400 bp, amplified with standard Illumina TruSeq PCR primers. Libraries were sequenced on the OSU CGRB Illumina HiSeq2500 machine.

Adapter sequences were removed from ChIP-seq reads and quality scores converted to Sanger format as described in [32]. Reads were aligned to the genome of Fol4287 using ‘bwa aln’ [88]. Duplicate reads were removed with Picard tools version 1.134 (http://broadinstitute.github.io/picard, MarkDuplicates) (Table S2). We used RSEG [89] to predict domains of enrichment using default settings and providing a bedfile with gaps in the assembly (N’s in the DNA sequence) as ‘deadzones’. We obtained two biological replicates for each experiment, one with relatively low and one with relatively high sequencing depth (Table S2). We estimate the reproducibility of our data by calculating the correlation of average read densities in 10 kb non-overlapping sliding windows. We obtained moderate correlation coefficients, ranging from 0,59 to 0,66 (Kendall’s Tau, Bonferroni-corrected P-value < 0.001) for H3K27me3 and H3K4me2 respectively. Reads have been uploaded to GEO (accession GSE121283).

### Synonymous and nonsynonymous substitutions

For Fol4287 genes based on alignments and trees with homologs in eight different *Fusarium* species, others [24] calculated omega values (d_N_/d_S_ ratios) and P-values indicating whether a model including site-specific selection fits the data better than a neutral model, for two different assumed distributions of d_N_/d_S_ ratios: one in which omega is either 0 or 1 (M1M2 in Fig 6), and one in which omega follows a beta distribution between 0 and 1 (M7M8 in Fig 6), where models with sites under positive selection (M2 and M8 respectively) have an additional category omega > 1. We downloaded these data from their Supplemental Online Material. In addition, we aligned Fol4287 protein sequences with their bidirectional best blast hits in the *F. verticillioides* proteome with Clustal Omega, used PAL2NAL to infer codon alignments and codeml to calculate d_N_ and d_S_ values based on these. We filtered out all genes with d_N_ or d_S_ > 1 (i.e. genes with multiple substitutions).

### SNP calling

Strains that belong to VCG030 are Fol002, Fol004, Fol007, Fol014, Fol018, Fol026, Fol029, Fol038, Fol073 and Fol074. The ‘clade III’- set consists of all strains in VCG030 and Fol016, Fol096, Fol072, Fol075, FolMN25, FoMN14, Fom001, Forc016, Forc024 and Forc031. Fol-CL25 also belongs to this clade, but no raw sequencing reads were publicly available for this strain. Raw sequencing reads were filtered and trimmed using fastq-mcf with ‘-q 20’ and a fasta file with Illumina adapter sequences. Trimmed reads were mapped to the genome sequence of Fol4287 using bowtie2 with ‘‐‐end-to-end ‐‐fr –reorder’ and ‘-I 100 -X 300’ for strains with an estimated fragment length of 200 (all strains in VCG030 except Fol007) and ‘-I 400 -X 600’ for other strains in clade III ((based on an estimated fragment length of 500). Bam files were processed individually for each sample. The files were sorted and duplicate reads were removed using Picard tools version 1.134 (http://broadinstitute.github.io/picard, MarkDuplicates). Variant calling was done with the Genome Analysis ToolKit [90] using Haplotypecaller with ‘-ploidy 1’. Indels were masked and SNP were sorted and filtered using VariantFilter with ‘maskExtension 5’, ‘clusterSize 3’, ‘clusterWindowSize 10’, ‘MQRankSum < −17.5’, ‘QD < 1’, ‘ReadPosRankSum < −8.0’, ‘FS > 60.0’, ‘MQ < 30.0’, ‘SOR > 7’, ‘MQ0 > 3’, ‘QUAL < 30.0’, ‘DP > 400’, ‘DP < 10’.

### Availability of data and scripts

To facilitate reproducibility of this work and allow others to perform similar analyses, the Python code and Ipython notebooks used to generate the Figures and to calculate statistics in this manuscript and its supplemental online material are available under GNU public license and can be downloaded from https://github.com/LikeFokkens/FOSC_multi-speed-genome. Reads from ChIP-seq experiments and the DNA sequencing of the two horizontal chromosome transfer strains (Fo47-1A and Fo47-2A, Fig 1B) have been submitted to the GEO (accession GSE121283) and the ENA (accession PRJEB29294) respectively.

## Supporting information captions

**Figure S1. Whole genome alignment of Fol4287 with Fom001.**

The whole genome alignment of tomato-infecting strain Fol4287 with melon-infecting strain Fom001 is represented here in a dotplot. Lines are coloured according to percent identity of the aligned segments, only alignments that are more than 90% identical and span more than 1 kb are included. Thin black vertical lines indicate Fol4287 chromosomes, grey, semi-transparent lines indicate scaffolds. Unpositioned scaffolds in Fol4287 are not included. The Fol4287 core chromosomes (1,2,4,5,7,8,9-13) are almost 99-100% identical in Fom001. With the exception of one translocation between chromosome 2 and chromosome 10, synteny is conserved in core chromosomes. In contrast, the accessory chromosomes are mostly absent. We expect that differences in host preference are largely determined by these accessory chromosomes. Chromosomal regions 1b, part of chromosome 15 and especially 2b are absent in sister species *F. verticillioides*, but largely present in Fom001 with high levels of sequence similarity. In contrast, chromsome 3, 6 and 14 are absent in Fom001.

**Figure S2. Presence/absence -coloured by % identity- of Fol4287 chromosome sequences in the genomes of 58 other *Fo* strains.**

The phylogenetic tree on the left was inferred using maximum likelihood from a concatenated alignment of 1194 core genes as described in van Dam et al. (Env. Microb. 2016). Leaf nodes are colored according to forma specialis; red: tomato, light-green: melon, green: cucumber, grey: cotton, petrol: watermelon, yellow: banana, purple: brassicaceae, brown: pea. Leaf nodes are shaped according to disease symptoms; round: wilting, square: root rot. Forc strains (green squares) are colored green but are pathogenic on cucumber, melon and watermelon. Strains that are not pathogenic on plants (to our best knowledge) have no shape, Fo47 and FoMN14 are non-pathogenic, FOSC-3a is pathogenic on immunocompromised humans. This plot indicates only presence of sequences and are not informative on synteny (see Figure S3 for plots colored according to alignment length): this plot only shows which segments are present, not whether they also accur on the same configuration/on the same chromosome. In Figure S2A and S2B, only alignments that span at least 1000 bp in the query genome and are at least 90% identical are included.

A. Comparison of core chromosomes, i.e. chromosomes that are largely present with high sequence similarity (>97% identity) in all strains. Part of chromosome 1 and chromosome 2 are considered accessory regions. Levels of sequence similarity correspond to the phylogeny. The level of sequence similarity drops in the subtelomeric regions. Chromosome 4 has a region in the middle with lower levels of sequence similarity, this region corresponds to the breakpoint of a chromosomal rearrangement when compared to *F. verticillioides*.

B. Comparison of fast-core and accessory chromosomes. The three smallest core chromosomes differ markedly from the other 8 core chromosomes depicted in Figure S2A. The percent identity in the alignments is lower, the difference is small but very consistent. Moreover, we observed more large-scale deletions in these chromosomes. The accessory chromosomes are present in only in a small subset of genomes. Some sequences are present with low sequence similarity (∼90%, depicted in blue), these could correspond to transposons. Alignments of chromosomal regions 1b and 2b and chromosome 15 with the Fom001 genome have higher sequence identity and synteny than of chromosome 3 and 6 (Figure 1, Figure S1). Moreover, chromosome 3 and 6 both contain a region that has been lost in strains Fol029, Fol018, Fol074 and Fol038. Chromosome 15 and chromosomal region 1b are absent Fol014, Fol029 and Fol018 and part of chromosome 14 has been lost in Fol018. Chromosome 1b, 2b, 3, 6, 15 and many of the unpositioned scaffolds hardly occur outside the clonal lineage of Fol4287. Some of the unpositioned scaffolds have a similar color pattern as core chromosomes, suggesting they are part of core chromosomes but could not be placed there due to lack of resolution in the optical map that was constructed for the reference assembly of Fol4287. Chromosome 14 and part of chromosome 3 and 6 clearly show hallmarks of large-scale horizontal transfer but there are also indications of smaller scale putative transfer events, e.g. part of chromosome 1b that is present in brassicaceae-infecting strains (indicated with purple circles in the phylogenetic tree). Part of chromosome 15 is also present in other strains, but with normal sequence similarity, suggesting that this region is lost in many strains rather than horizontally transferred. Comparisons to outgroup species are needed to confirm this.

C. Presence-absence of core, fast-core and accessory chromosomes with more lenient cutoffs: all alignments that are more than 80% identical in sequence and span more than 100 basepairs are included. The patterns we observed above are not dependent on specific cut-offs, as we observe then with more lenient cut-offs as well.

**Figure S3. Presence/absence -coloured by synteny- of Fol4287 genome sequences in the genomes of 58 other *Fo* isolates.**

The phylogenetic tree on the left was inferred using maximum likelihood from a concatenated alignment of 1194 core genes as described in van Dam et al. (Env. Microb. 2016). Leaf nodes are colored according to forma specialis; red: tomato, light-green: melon, green: cucumber, grey: cotton, petrol: watermelon, yellow: banana, purple: brassicaceae, brown: pea. Leaf nodes are shaped according to disease symptoms; round: wilting, square: root rot. Forc isolates (green squares) are colored green but are pathogenic on cucumebr, melon and watermelon. Isolates that are not pathogenic on plants (to our best knowledge) have no shape, Fo47 and FoMN14 are non-pathogenic, FOSC-3a is pathogenic on immunocompromised humansIn Figure S3A and S3B, only alignments that span at least 1000 bp in the query genome and are at least 90% identical are included.

A. Comparison of core chromosomes. Part of chromosome 1 and chromosome 2 are considered accessory regions. Within the clonal line of Fol4287 and with Fom001, alignments can span as long as and longer than 50 Mb (plotted in yellow). As with sequence similarity, synteny levels drop at the subtelomeric regions and in the middle region of chromosome 4 that corresponds to the breakpoint of a chromosomal rearrangement when compared to *F. verticillioides*. Fol4287 self-alignment does not produce only alignments of > 50 Mb in length because the assembly contains gaps. Chromosomal region 2B is relatively fragmented (contains > 2 times the number of gaps as e.g. chromosomal region 1B, or Supercontig_2.25 that is of comparable size and is located on chromosome 3) which is reflected in the low level of synteny in the Fol4287 self-alignment.

B. Comparison of fast-core and accessory chromosomal regions. The three smallest core chromosomes differ from the other 8 core chromosomes depicted in Figure S3A, synteny conservation is less conserved in more distant isolates. Part of chromosome 13 is not syntenic to Fom001, it aligns to a number of relatively small contigs. Notably, this region is lost in Fom013. The accessory chromosomes are present in only in a small subset of chromosomes. While for chromosome 14 sequences are conserved in tomato-infecting isolate, synteny is not, in contrast to the region on chromosome 3 and 6 that is shared between tomato-infecting isolates, where we do observe conserved synteny.

C. Presence-absence of core, fast-core and accessory chromosomes with more lenient cutoffs: all alignments that are more than 80% identical in sequence and span more than 100 basepairs are included. The patterns we observed above are not dependent on specific cut-offs, as we observe then with more lenient cut-offs as well.

**Figure S4. Gene and repeat density**.

For 10 kb non-overlapping windows, we here plot the fraction of base pairs that is part of a gene (blue) or a predicted transposable element (grey). The accessory chromosomes and chromosomal regions are rich in repeats compared to the core chromosomes, including the fast-core chromosomes. Chromosomes are scaled according to size.

**Figure S5. Fast-core and accessory chromosomes have lower gene expression levels**.

The expression level in reads per kb per million mapped reads (RPKM), averaged over three biological replicates. Per chromosome, there are two panels: the top panel shows *in vitro* gene expression in black dots and the bottom panel *in planta* gene expression in blue dots. The x-axis show the position of the gene on the genome, the y-axis the RPKM. Note that the y-axis is log-scaled. We drew a red solid line at y = 5 for reference. On core chromosomes, gene expression levels are mostly above this line, where on fast-core and accessory chromosomes, most gene expression levels are under 5 RPKM.

**Figure S6. Fast-core and accessory chromosomes have relatively more differentially expressed genes than core chromosomes**.

The x-axis show the position of the gene on the genome, the y-axis the log2fold change in gene expression (*in planta/in vitro*) per chromosome. Zero average read counts have been replaced by 0.1 to avoid infinite ratio’s (as in Van der Does et al. PloS Genetics 2016). Differentially expressed genes are highlighted in blue squares. Not only fast-core chromosomes and chromosome 14 but also the central fast-core region on chromosome 4 and the subtelomeric regions have many differentially expressed genes.

**Figure S7. Functional categories that are overrepresented on the fast-core chromosomes**.

Fraction of annotated genes that are assigned to a FunCat category that is significantly overrepresented on the fast-core chromosome, compared to all genes on the genome (P-value < 0.001 after FDR correction). Categories that are also overrepresented when comparing up-regulated genes to the background of all fast-core genes are in bold. Percentages are defined with respect to the total number of annotated genes.

**Figure S8. Read density from ChIP-seq experiments targeted at H3K27me3-ated or H3K4me2-ated regions**.

Average read densities in 10.000 bp non-overlapping sliding windows are depicted in green (H3K4me2) and orange (H3K27me3). Read densities are normalized with respect to RPKM to allow for comparisons across experiments. Chromosomes are scaled to the length of their sequence in the assembly. The core chromosomes - except chromosome 11, 12 and 13- are largely enriched for H3K4me2, except at the subtelomeric regions. These small core chromosomes, designated fast-core chromosomes, are enriched in H3K27me3, similar to accessory chromosomes. Also the middle region in chromosome 4, which is also more divergent in terms of sequence similarity and synteny (Figure S2A, Figure S3A) is enriched in H3K27me3 as well. The two histone marks appear to be mutually exclusive.

A. Experiments with high sequencing depth: H3K4me2, experiment id 1358: 19735885 reads, of which 18129305 mapped (91,86 %); H3K27me3, experiment id 1360: 21467536 of which 16243857 mapped (75,76 %), see Table S2.

B. Replicate experiments, with lower sequencing depth: H3K4me2, experiment id 806: 9078423 reads, of which 6699238 mapped (73,79 %); H3K27me3, experiment id 808: 7891973 of which 5519933 mapped (69,94 %), see Table S2.

**Figure S9. Core chromosomes are enriched for H3K4me2 and fast-core and accessory chromosomes for H3K27me3**.

For each chromosome or chromosomal region (x-axis) we plot how much is covered in a DOE (top panels) and the size-distribution of DOEs (bottom panels, violinplots). Lineage-specific regions are denoted with a ^#^ on the x-axis. The regions on chromosome 3 and on chromosome 6 that are associated with pathogenicty on tomato are denoted with a ^P^. Bars with thick lines represent the coverage per chromosome category (core, fast-core, lineage specific or pathogenicity) rather than per chromosome.

A. Coverage and size distribution of DOE inferred based on two H3K4me2 experiments (1358 ‘H3K4me2 (a)’ and 806 ‘H3K4me2 (b)’. B. Coverage and size distribution of DOE inferred based on two H3K27me3 experiments (1360 ‘H3K27me3 (a)’ and 808 ‘H3K27me3 (b)). The coverage of DOEs is lower for 806 ‘H3K4me2 (b)’ and 808 ‘H3K27me3 (b), than for 1358 ‘H3K4me2 (a)’ and 1360 ‘H3K27me3 (a)’, but the distribution are not qualitatively different.

**Figure S10. Overlap of domains of enrichment with genes, promoters and repeats**.

We used RSEG to identify domains enriched in histone marks H3K4me2 (A) or H3K27me3 (B), based on mapped reads generated in two independent ChIP-seq experiments for each mark (left and right panels, 1358 and 806 for H3K4me2, 1360 and 808 for H3K27me3) (see Materials and Methods for more detail). For each chromosome or chromosomal region, we determine the percentage of genes (top panel, light blue), promoter regions (defined as up to 1000 base pairs upstream of first exon, second panel, dark blue), and repeats (as predicted by RepeatMasker, excluding low-complexity regions and simple repeats, third panel, grey) that overlap (for more than 90%) with a domain of enrichment. To compare whether certain elements (e.g. genes) overlap more than expected with a domain of enrichment, we also plot how much of a chromosome or chromosomal region is part of a domain of enrichment (bottom panels, dark grey on light grey background). Average values per chromosome category are also included in the histogram, and designated with black boxes.

A. H3K4me2 domains occur mainly on the core chromosomes, and very little on the fast-core and accessory chromosomes. Domains of H3K2me2 are relatively underrepresented in repeat regions.

B. H3K27me3 domains occur mainly on the fast-core and accessory chromosomes, and very little on the core chromosomes. Domains of H3K27me3 are overrepresented in repeat regions on the core chromosomes.

**Figure S11. dN, d_S_ and dN/dS values along the genome**.

The number of synonymous substitutions per synonymous site d_S_, the number of non-synonymous substitutions per nonsynonymous site d_N_ (top panels, d_S_ in blue, d_N_, in black traingles) and their ratio (bottom panel, grey dots) of genes are plotted on the genome. Genes with d_N_/d_S_ > 1 are highlighted with red squares. On the x-axis are chromosomes, dotted vertical lines denote supercontigs that comprise these chromosomes. Y-axes are in log scale. Subtelomeric regions have higher d_N_ and d_S_ values, while their ratio is consistent along the genome. The same holds true for the fast-core chromosomes and the middle region on chromosome 4 that is also enriched in H3K27me3 and depleted in H3K4me2. The accessory chromosomes have very little data points, as values are calculated with respect to bidirectional best blast hits in *F. verticillioides* and these chromosomes do not occur in this species.

**Figure S12. Frequency distribution of d_N_ and d_S_ of core and fast-core genes**.

We compare the distributions of all core (light-blue) or all fast-core (light-red) genes to two subsets of genes (dark-blue and dark-red). One subset consists of genes that are differentially expressed during infection (DEGs, left panels) and one of genes that are in an H3K27me3-enriched domain (H3K27me3, right panels).

A. Kernel density estimation of the frequency distribution of d_N_. Fast-core genes have higher d_N_ values than core genes. Fast-core DEGs have low values d_N_ compared to all fast-core genes, while for core genes there is no difference (left panel). Core genes that are in an H3K27me3-enriched domain have the same distribution as fastcore genes.

B. Kernel density estimation of the frequency distribution of d_S_. Fast-core genes have higher d_S_ values than core genes. Core DEGs have higher d_S_ values than other core genes, for the fast-core the distribution of d_S_ of DEGs does not differ much from that of all fast-core genes. Again, core genes that are in an H3K27me3-enriched domain have the same distribution as fast-core genes.

**Figure S13. SNP density per 10 kb sliding window, per chromosome**.

We here plot the cumulative SNP density in 10000 bp, non-overlapping sliding windows. Per chromosome there are two panels: the top panel shows the cumulative SNP density for all strains in Clade III (see tree in the legend) in purple, whereas the bottom panel shows shows the cumulative SNP density for all strains in Vegetative Compatibility Group 030 (VCG030, dark-grey).

**Figure S14. SNP density per 10 kb sliding window, per chromosome**.

We here plot the cumulative SNP density in 10000 bp, non-overlapping sliding windows. Per chromosome there are two panels: the top panel shows the cumulative SNP density for all strains in Vegetative Compatibility Group 030 (VCG030, dark-grey), using a different y-axis scale than in Figure S13 to show that the distribution of SNPs on fast-core chromosomes is not different from that on core chromosomes.

**Table S1. Strains used in this study**.

**Table S2. Gene expression levels and fold change in gene expression per gene**

**Table S3. Functional analysis of differentially regulated genes on pathogenicity region on chromosome 3 and 6**.

**Table S4. Functional analysis of differentially regulated genes on chromosome 14**.

**Table S5A. Functional enrichment of genes that reside on chromosome 11, 12 and 13**.

**Table S5B. Functional enrichment of up-regulatedgenes that reside on chromosome 11, 12 and 13**.

**Table S6. Sequencing depth, mapping and putative PCR duplicates in the ChIP-seq experiments**.

**Table S7A. Domains of enrichment of H3K4me2**.

**Table S7B. Domains of enrichment of H3K27me3**.

**Table S8. Statistics of selection per gene**.

